# Social organization and biting performance in *Fukomys* mole-rats (Bathyergidae, Rodentia)

**DOI:** 10.1101/325720

**Authors:** P.A.A.G. Van Daele, N. Desmet, D. Adriaens

**Keywords:** caste, eusociality

## Abstract

A caste system, based on work activity and reproduction, has been proposed in the two African mole-rat species which are generally considered eusocial, *Heterocephalus* glaber and *Fukomys damarensis*. Social behaviour in other *Fukomys* species is key to understanding evolution of sociality within bathyergids, which display a social continuum among species from solitary to eusocial. Furthermore, insight in the social structure of colonies may be instrumental in understanding the observed, extensive intraspecific morphological variation and ultimately help species delimitation. For the first, time social organisation was studied in a colony of wild-caught *Fukomys micklemi* (Sekute cytotype) from Zambia. Data were collected on work behaviour and analysed against morphological variables and biting performance. Although there was considerable variation in the amount of work performed by each individual, clearly distinguishable castes were not found. Castes might represent an artificial subdivision, which does not necessarily reflect the dynamic changes within a colony. Variation in work may be the result of an ongoing process of continuous change, whereby a colony undergoes a certain evolution that is reflected in developmental patterns of individuals. Ecological factors will undoubtedly play an important role in colony evolution. Consequently, predictions made according to the Aridity Food Distribution Hypothesis should be tested, taking into account the possibility of a dynamic model as described here. The relation between biting performance and behavioural traits was investigated for the first time. Whereas differences in biting performance were strongly correlated with morphological parameters, relation between work and biting performance remains unclear.

## Introduction

Ever since the naked mole-rat (*Heterocephalus glaber*) and the Damaraland mole-rat *Fukomys damarensis* were found to fit the criteria for eusociality in insects, the definition of eusociality has been debated (Bennett and Faulkes 2000, Burda et al. 2000). While some authors promote a strict application of the term eusocial (Crespi and Yanega, 1995), others suggest the use of a continuum (Gadagkar, 1994; Keller and Perrin, 1995; Sherman et al., 1995). According to Burda et al. (2000) mammalian eusociality is characterized by the following criteria: (1) reproductive altruism with a reproductive division of labour and cooperative alloparental brood care, (2) an overlap of generations and (3) permanent philopatry. Whereas the existence of different worker castes is a typical feature of eusociality in insects, it is not a paramount criterion in this definition. Nevertheless, the existence of different castes is often seen as an important feature defining eusociality in African mole-rats (Bennett and Faulkes, 2000). The eusocial mole-rat species are characterized by colonies with a caste system. Three castes have been recognized: a reproductive caste, in which one female and up to three males reproduce, and among non-reproducing helpers two worker castes, viz. a frequent and infrequent worker caste (Bennett and Jarvis 1988, Bennett 1989, 1990, Jacobs et al. 1991, Gaylard et al. 1998, Wallace and Bennett 1998, Scantlebury et al. 2006). Social organisation in other *Fukomys* species has not been studied in detail, but most species are assumed to occupy an intermediate position (cf. Faulkes and Bennett 2007 and references there-in).

The Aridity Food Distribution Hypothesis (AFDH) offers a framework for the evolution of sociality in rodents, and in mole-rats (Bathyergidae, Rodentia) in particular (Lovegrove, 1991; Jarvis et al., 1994; Faulkes et al., 1997b; Bennett and Faulkes, 2000; Faulkes and Bennett, 2007). According to the AFDH sociality and cooperative behaviour have evolved as an adaptive response to an unpredictable environment, whereby an unpredictable environment is defined as an environment with scattered food resources and low and variable rainfall. Since burrowing is an energetically costly activity, with the costs increasing dramatically with soil hardness, most of the burrowing will be carried out when the soil is moist and soft (Jarvis et al., 1998). Consequently, when rainfall is both unpredictable and low, the opportunities for digging new tunnels to search for food or disperse and form new colonies are limited. Furthermore, plants occurring in these habitats are adapted to arid conditions. Adaptations include the formation of swollen roots and tubers in which reserves are stored. Vegetative reproduction of these plants implies a patchy and/or widely dispersed distribution of clumps compared to plants occurring in mesic regions. While the swollen roots and tubers of these plants form an excellent source of food and water for mole-rats, the patchy distribution increases the risk of unsuccessful foraging (Faulkes and Bennett, 2007). The AFDH predicts that living in social groups can decrease the risk of unsuccessful foraging for food resources. Whereas the chance of a single mole-rat finding a clump of roots through energetically costly burrowing are low, a group of animals will have a significantly higher chance of discovering such a food source. Once a patch of swollen tubers is found, it will provide food and moisture for a considerable time (Jarvis et al., 1994). Thus, in conditions where chances of dispersal and successful establishment of a new colony are slim, a division of labour is likely to occur, eventually leading to a caste system in arid habitats (Jarvis and Bennett, 1991).

It should, however, be noted that the outcomes of the various studies are not equivocal, neither is the reinterpretation of the results by various authors (cf. e.g. Burda et al. 2000). Evolution of bathyergid sociality from a solitary ancestor was supposed to be possible thanks to the body mass independent metabolism, which was understood as an exaptation for the evolution of sociality in African mole-rats (Lovegrove and Wissel 1988, Lovegrove 1991). Following this assumption, groups i social species would benefit from a large workforce of small foragers, with no increase of metabolic costs. However, the meta-analysis of representative data on resting metabolic rates in 14 bathyergid taxa using the method of phylogenetic contrasts clearly demonstrated that resting metabolic rates in African mole-rats follow a typical mammalian body pattern (Zelová et al. 2010).

Another challenge for the AFDH stems from the fact that not biomass or abundance of the food supply is important but also its spatial distribution (Jarvis et al. 1994, 1998). In spite of the fact that food resources in mesic areas were considered to be randomly or regularly distributed, analysis of food supply from different natural mesic habitats (grassland, miombo woodland) showed a clumped distribution of food resources I the solitary silvery mole-rats (Lövy et al. 2012). This finding falsifies the prediction of the AFDH that solitary species would avoid these areas.

*Fukomys* mole-rats are obligatory subterranean, living in a complex tunnel system which they dig out using their incisors (chisel-tooth digging). Furthermore, the biting apparatus is used for feeding on hard geophytes, defending the colony and during social interactions (Bennett and Faulkes, 2000). The biting apparatus thus plays an important role in the ecology of mole-rats and shows specific morphological adaptations to their subterranean lifestyle, such as hypertrophied jaw muscles and lips closing behind the incisors. Although ecological factors may have substantial consequences on skull morphology and biting performance (Herrel et al., 2006), phylogenetic and mechanical constraints will pose limits on the evolution of morphology (Irschick et al., 1997; Herrel et al., 2004; Kohlsdorf et al., 2008). Vanhooydonck et al. (2001) argue that evolutionary trade-offs may constrain the phenotypic radiation of a group. When this is applied to mole-rats, severe constraints on morphological variation could be expected on the basis of their subterranean lifestyle. Indeed, species within the genus *Fukomys* show high morphological similarity, despite impressive variation at the chromosomal and molecular level. (Faulkes et al., 2004; Ingram et al., 2004; Van Daele et al., 2004, 2007a). Within the genus, different clades can be distinguished based on DNA markers. Taxa within all of these clades show a considerable amount of DNA sequence and/or chromosomal and/or morphological variation. Species boundaries therefore remain unclear (Van Daele et al., 2004, 2007b). One way of gaining insight in the morphological evolution of mole-rats is through studies of morphological variation at the colony level.

In spite of the apparent similarity in external morphology between species within *Fukomys*, preliminary studies on the interspecific level show the existence of considerable shape and size polymorphism (with different cranial and mandibular morphotypes) among closely related chromosomal races (Van Daele et al., 2006; Murtas et al., 2007; Addendum 1 in Van Daele 2012). Additionally a recent comparative study among three species of *Fukomys* has shown interspecific differences in biting performance, which may be related to skull anatomy and skull shape (Van Daele et al, 2009). Furthermore, it is clear not only intraspecific variation in skull morphology but also in biting performance is larger than the interspecific variation. This raises the question if the high amount of intraspecific variation could be the result of caste-related differences in biting performance. In this respect, it could be hypothesized that frequent workers bite harder than infrequent workers and would show related differences in skull shape and biting performance. The possible existence of different worker castes has not been tested for *Fukomys micklemi*. Behavioural observations will therefore provide insight into the social structure of this species and may therefore help to clarify evolution of (eu)sociality in African mole-rats. An understanding of variation in behaviour and biting performance may contribute to a better understanding of morphological evolution of African mole-rats and of a possible mechanism behind caste formation. Combined with molecular genetic data, this may ultimately aid in clarifying species boundaries within *Fukomys*.

We carried out behavioural observations in order to map out variation in the type and amount of work performed by individual mole-rats. Behavioural data were linked to bite force measurements, resulting in the mapping of intraspecific variation in biting performance at the colony level. In order to get a complete view of biting performance related to castes and morphology, the relationship between bite force and quantitative aspects of morphology (mass and skull measurements) was determined. This allowed to test whether or not the intracolonial morphological differences (Van Daele et al., 2006) may be adaptive, i.e. if they are the result of differential selection pressure on different castes, whereby a higher performance pressure in a frequent worker caste is expected.

## Materials and methods

### Specimens

The studied specimens belong to a single colony of *Fukomys micklemi*, more specifically of the Sekute cytotype, 2n = 56 (Van Daele et al. 2004). Animals were kept in a vivarium at Ghent University. For husbandry details see Desmet et al. 2014; Van Daele 2012). *Fukomys micklemi* is closely related to the intensively studied *Fukomys damarensis* of the chromosomally hyper-diverse central Zambezian lineage (*F. micklemi* clade, 2n = 42-68; Van Daele et al. 2007a, b). All specimens in this study were wild-caught in Southern Zambia (Sekute area, 17°39’S, 25°37’E) in September 2008, using traditional methods (Jarvis 1991a, b). For an overview of the composition of this colony see Table 9.1. During the observation period no breeding occurred in this colony.

### Behavioural observations

Different methodological alternatives were evaluated in a preliminary study. It was concluded that a combination of behavioural sampling with continuous recording and a terrarium set up with a digging unit were the most efficient approach, providing the best behavioural resolution (for details see Desmet et al. 2014.; Van Daele 2012). All three categories of work behaviour(Addendum 2 in Van Daele 2012) were included: digging (including backshoveling, foreleg digging, sweeping using hindfeet and chisel tooth digging), transporting (including carrying food, carrying nest material, dragging, sweeping and kicking backwards) and allogrooming (non-genital and genital allogrooming).

### Bite force

In vivo bite forces were measured using an isometric Kistler force transducer, which was connected to a charge amplifier (adapted from Herrel et al., 1999). The animals were also weighed to the nearest gram. Animals were taken out of their terraria and placed individually in a small plastic cage. When the animals were approached, this usually resulted in a characteristic defensive threat posture. Most animals would immediately bite any approaching object. Other animals needed to be spurred on by blowing at them. The animals were induced to bite at least five times. The highest bite-force readout was used as an estimator of maximal biting performance. Head length and head height were measured to the nearest 0,1 millimetre on pictures of standardized lateral views, using ImageJ 1.41o software (Schneider et al. 2012).

### Analysis

For each individual, the total time spent out of the nest was calculated, as well as the time spent digging, time spent transporting and the time allogrooming. Statistical analyses were conducted in Statistica V 7.0 (Statsoft, 2004).

## Results

The study colony has an unusual structure. Whereas a typical colony of *Fukomys* mole-rats contains one reproductive female and one to three reproductive males (Jarvis et al., 1994, own observations in wild-caught and captive colonies), the studied colony contains two breeding females (ID 34 and ID 62). Both females had visible mammae and a perforated vagina. Furthermore, both females were seen copulating (both with the ID 61 male) during observations. One of the females (ID 62) had a small pup which could be assigned to her with great certainty. Both this female and the ID61 male were very protective over it. They typically positioned themselves in such a manner that the pup was placed between them and they immediately responded to disturbance with an aggressive defensive display.

### Work behaviour

Although there is considerable variation in the amount of work performed by each individual no clear indication of the existence of worker castes can be discovered (Table 10.1, Fig. 10.1). The total amount of work per individual (expressed as percentage of the total work performed by the entire colony) ranges from 1.37% (ID 62) to 8.39% (ID 28). The distribution between these values, however, is continuous. Similar results were found for digging and transporting (not shown).

**Fig. 10.1.**
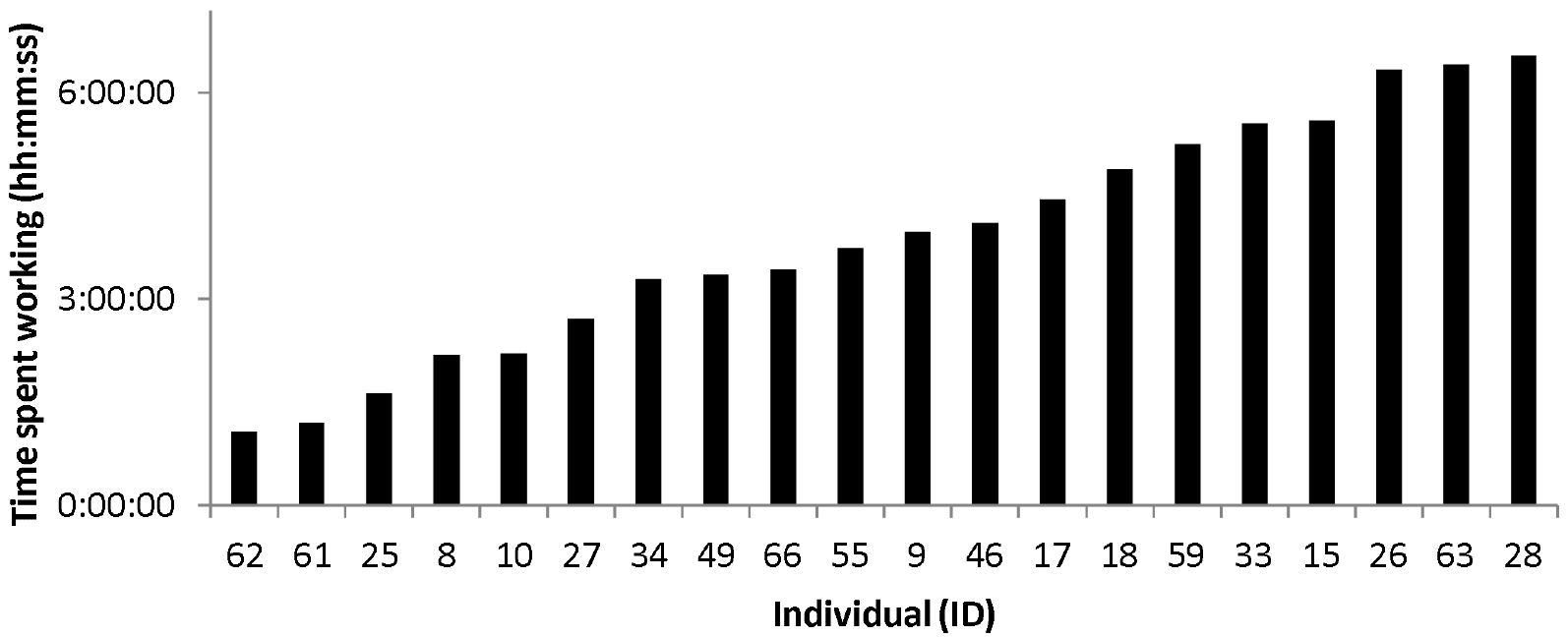
Total work performed by each individual of a colony of *F. micklemi*.

Allogrooming is mostly restricted to a few individuals, with one individual (ID 33) doing almost one third of the grooming. This individual also spent a lot of time in the nest. Furthermore, there is variation in the amount of time one individual would spend performing a particular kind of work (Fig. 10.2). Some individuals will spend more time transporting (e.g. ID 34), while the primary activity of others will be digging (e.g. ID 27 and ID 61). Consequently, although there is no clear distinct frequent worker caste, different individuals will be primarily involved in different types of work.

Males and females did not differ significantly in the amount of work done ((t_df 19_ p =0.48; mean (females) = three hours 54 minutes; mean (males) = three hours 52 minutes)). Reproductive female 62 and reproductive male 61 were the two animals which performed the least amount of work (1.37% and 1.54% respectively). ID 34 accounted for 4.21% of the work (the largest part of this was transporting nest material; accounting for 8.49% of total transporting behaviour by the colony)

There was no clear correlation between the amount of work and body mass (r= −0.39, p=0.88) or head height (r= −.040, p=0.88), but head length was negatively correlated with work (r= −0.75, p=0.0002) (Fig.10. 3). Time spent working was poorly correlated with active time (i.e. time spent out of the nest; r= 0.48; p=0.02; Fig. 10.4).

**Table 10.1.** Work behaviour in *F. micklemi* Amount of work (per behaviour and total) performed by each individual (expressed as a percentage of the total for the entire colony).

| Individual (ID) | Sex | Digging (%) | Transporting (%) | Allogrooming (%) | Total (%) |
| --- | --- | --- | --- | --- | --- |
| 8 | F | 3.34 | 2.34 | 1.28 | 2.81 |
| 9 | F | 4.42 | 5.27 | 11.41 | 5.09 |
| 10 | F | 1.72 | 4.12 | 2.99 | 2.84 |
| 15 | F | 9.07 | 5.36 | 3.26 | 7.18 |
| 17 | M | 5.73 | 6.22 | 0.00 | 5.71 |
| 18 | F | 8.01 | 4.32 | 5.80 | 6.28 |
| 25 | M | 2.93 | 1.31 | 0.27 | 2.09 |
| 26 | M | 10.31 | 6.12 | 2.69 | 8.13 |
| 27 | F | 5.12 | 1.81 | 0.98 | 3.48 |
| 28 | F | 8.54 | 7.89 | 11.85 | 8.39 |
| 33 | F | 7.47 | 4.52 | 30.39 | 7.13 |
| 34 | BF | 0.83 | 8.49 | 0.47 | 4.21 |
| 46 | F | 5.18 | 5.63 | 2.62 | 5.27 |
| 49 | M | 5.15 | 3.69 | 0.46 | 4.31 |
| 55 | M | 5.19 | 4.67 | 1.50 | 4.80 |
| 59 | F | 3.81 | 10.01 | 8.14 | 6.74 |
| 61 | M | 2.25 | 0.85 | 0.00 | 1.54 |
| 62 | BF | 0.96 | 1.95 | 0.20 | 1.37 |
| 63 | M | 6.01 | 11.10 | 4.98 | 8.22 |
| 66 | F | 3.95 | 4.33 | 10.70 | 4.40 |
| **TOTAL** |  | 100 | 100 | 100 | 100 |
| **Legend** | M: male | F: female | BF: breeding female |  |  |

**Fig. 10.2.**
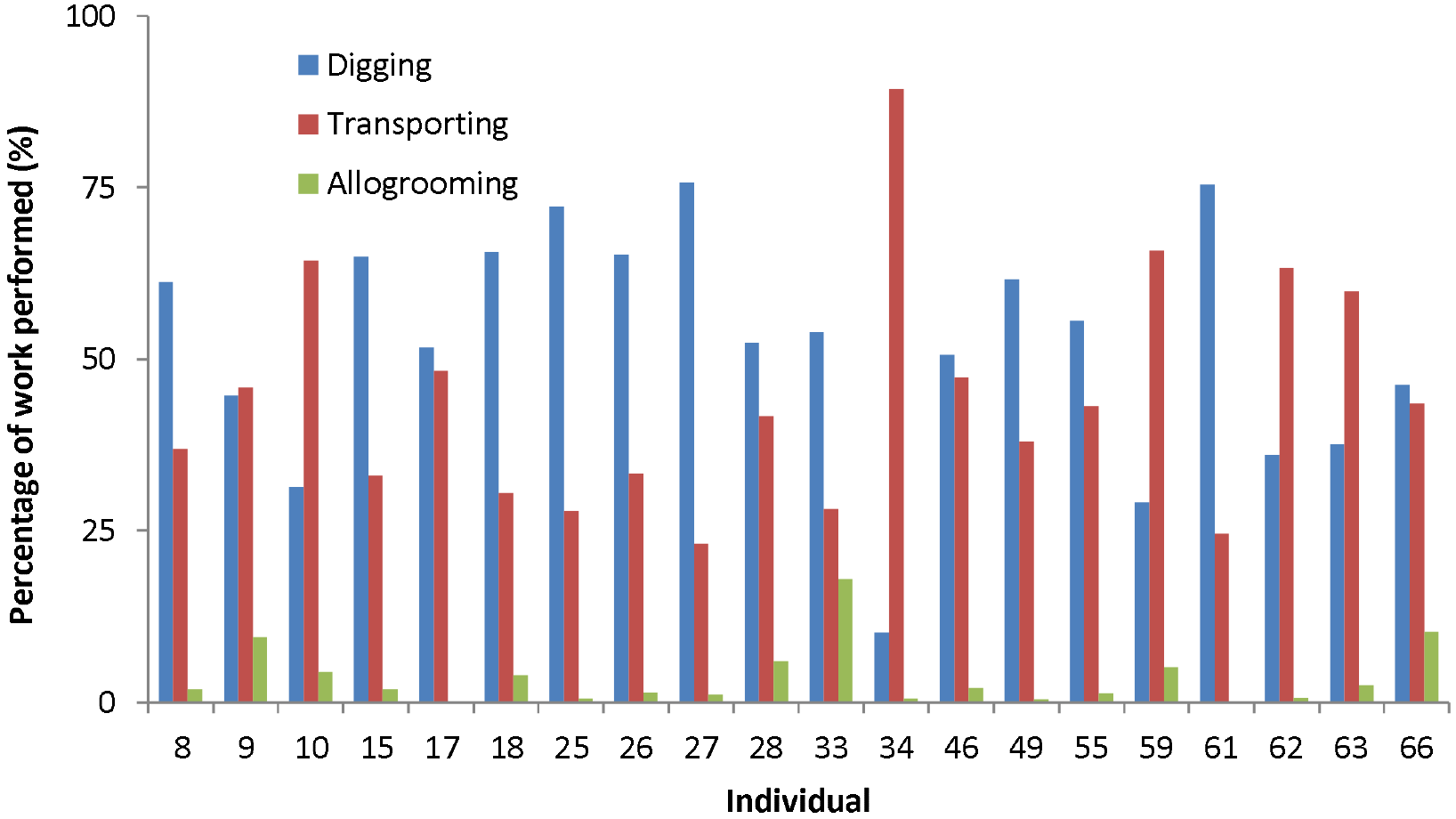
Work behaviours in *F. micklemi*: Each category of work expressed as the percentage of the total work performed by that individual.

**Fig. 10.3.**
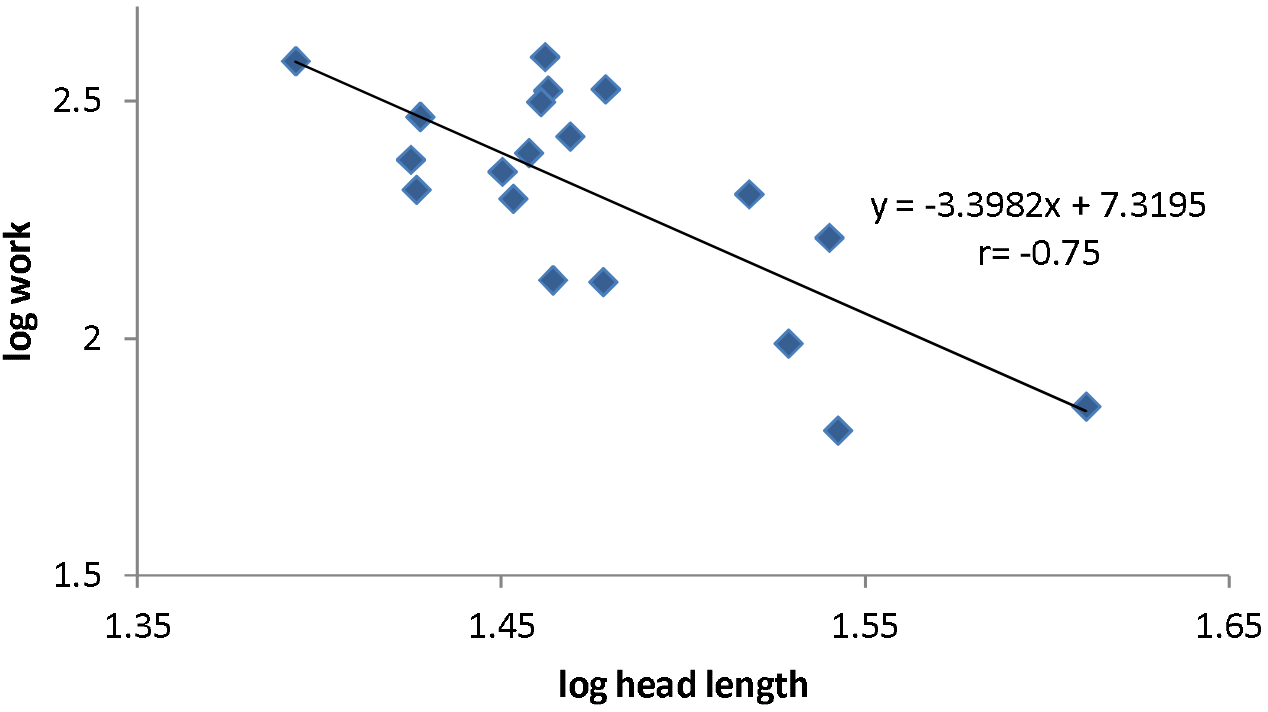
Relationship between work and head length in a *Fukomys micklemi* colony.

**Fig. 10.4.**
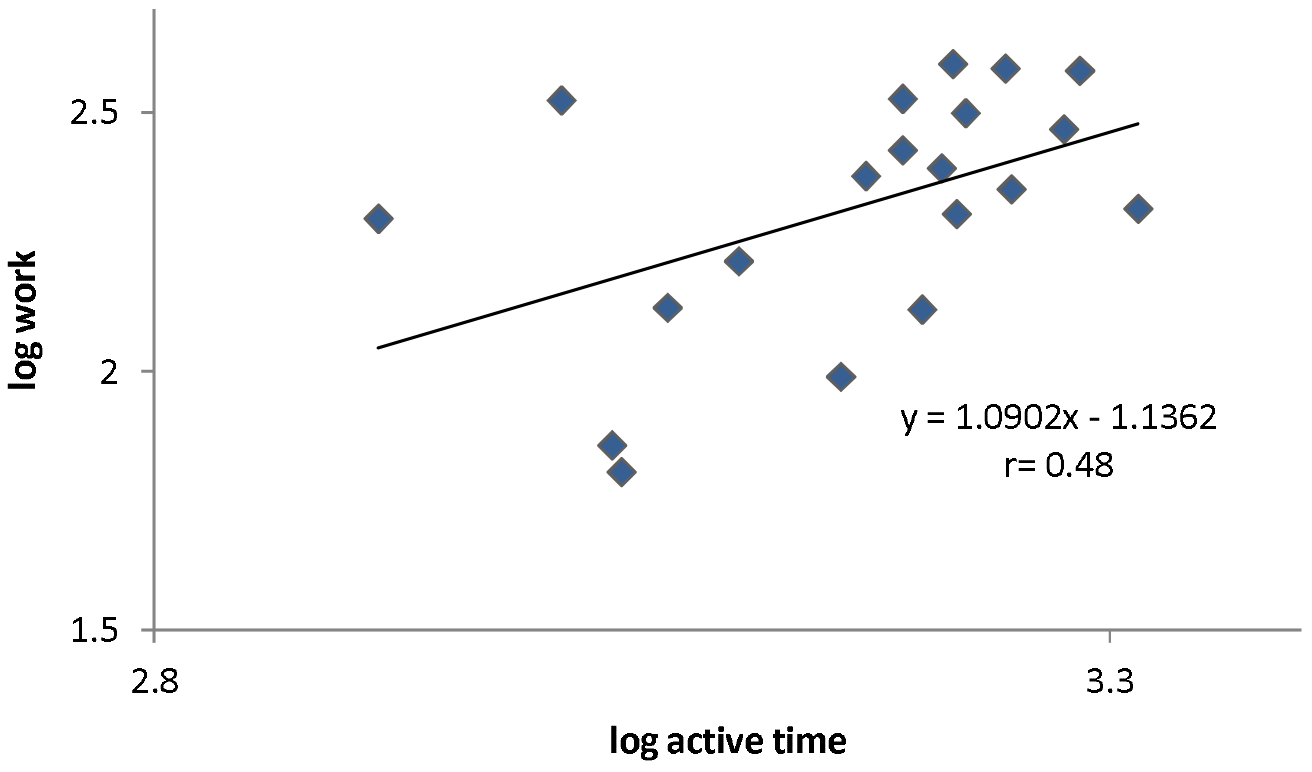
Relationship between active time and work in a *Fukomys micklemi* colony.

### Bite force

#### Field versus captivity

The animals of the study colony showed no significant difference between maximum bite force in the field compared with maximum bite force measured in captivity (see Table 10.2; t-test: p=0.61). When body mass-corrected bite force was compared, however, there was a significant decrease in bite force per unit of mass in captivity in comparison with field measurements (t_df 19_ p<0.001; Fig. 10.5). This could be explained by a significant increase in overall mass in captivity (t-test: p<0.001). No significant differences were found between male and female bite force (t_df 19_ p=0.30 and p=0.27, respectively for measurements in the field and in captivity; Fig. 10.5).

**Fig. 10.5.**
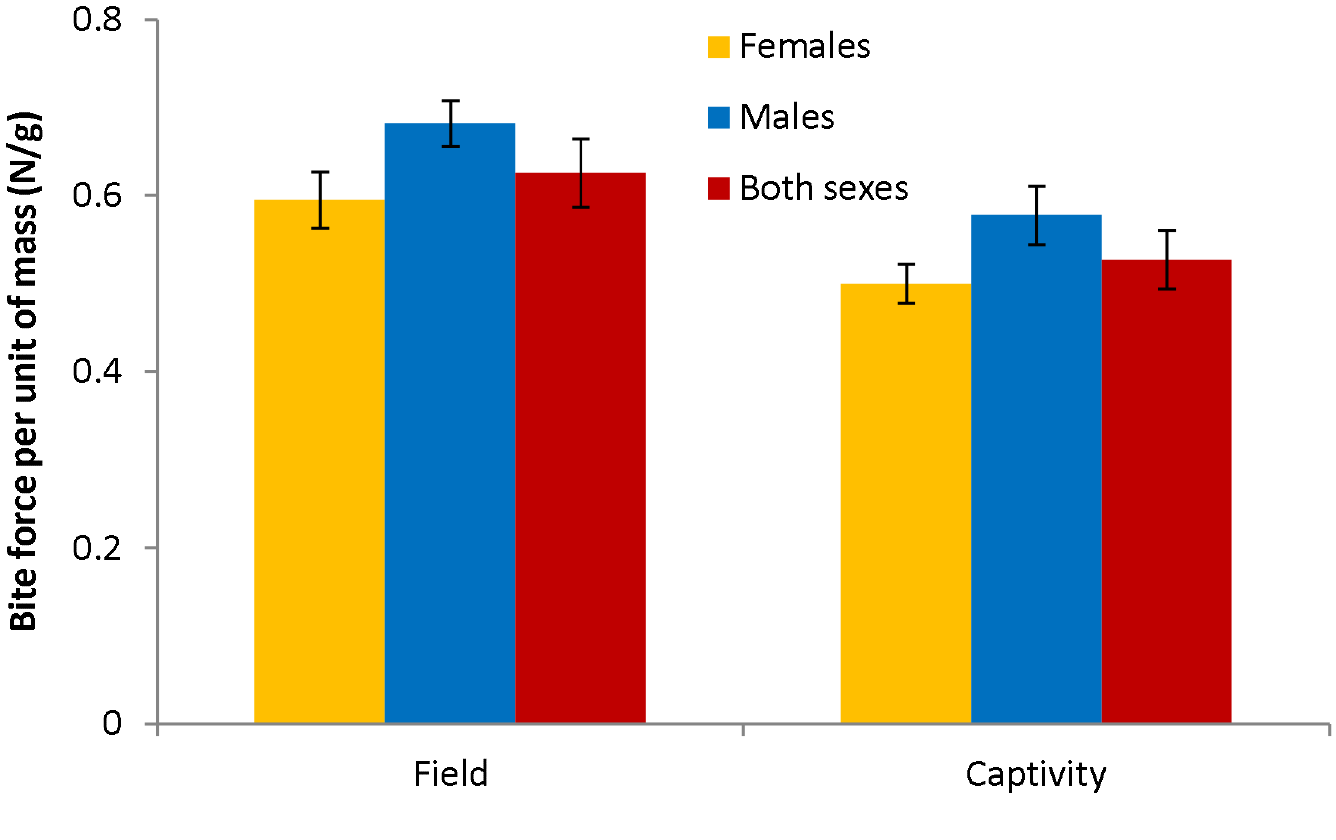
Comparison of bite force per unit of mass for measurements in the field and in captivity in a colony of *Fukomys micklemi*.

**Table 10.2.** Mass, bite force and head dimensions for a colony of *F. micklemi*

| Individual (ID) | Sex | M1 (N) | BF1 (g) | M2 (g) | BF2 (N) | HL (mm) | HH (mm) |
| --- | --- | --- | --- | --- | --- | --- | --- |
| 8 | F | 44 | 23 | 60 | 23 | 30,1 | 28,6 |
| 9 | F | 72 | 36 | 82 | 40 | 26,6 | 34,0 |
| 10 | F | 77 | 36 | 80 | not motivated | 29,1 | 34,4 |
| 15 | F | 75 | 41 | 90 | 51 | 30,1 | 32,2 |
| 17 | M | 114 | 89 | 132 | 96 | 29,4 | 37,2 |
| 18 | F | 55 | 6 | 73 | 13 | 26,8 | 31,6 |
| 25 | M | 90 | 64 | 104 | 75 | 33,8 | 36,5 |
| 26* | M | 63 | 38 | 92 | 45 | - | - |
| 27 | F | 78 | 49 | 96 | 39 | 34,7 | 34,0 |
| 28 | F | 80 | 46 | 91 | 44 | 29,0 | 34,7 |
| 33 | F | 77 | 48 | 93 | 55 | 29,0 | 31,8 |
| 34 | BF | 104 | 64 | 108 | 69 | 28,4 | 39,8 |
| 46 | F | 76 | 53 | 89 | 45 | 28,7 | 35,3 |
| 49 | M | 102 | 85 | 106 | 70 | 33,0 | 37,1 |
| 55 | M | 46 | 26 | 65 | 20 | 28,2 | 33,8 |
| 59 | F | 72 | 45 | 101 | 54 | 28,9 | 32,9 |
| 61 | M | 124 | 106 | 151 | 115 | 40,8 | 40,8 |
| 62 | BF | 84 | 55 | 83 | 49 | 34,9 | 34,7 |
| 63 | M | 14 | 6 | 37 | 14 | 24,7 | 29,5 |
| 66 | F | 57 | 51 | 84 | 44 | 26,7 | 33,0 |
| Mean |  | 76,00 | 6,00 | 37,00 | 50,95 | 30,15 | 34,31 |
| SD |  | 25,39 | 25,23 | 24,38 | 26,25 | 3,78 | 3,10 |

**Legend** m: male f: female bf: breeding female
\* died after observations had been finished
M1: body mass in the field
M2: body mass in captivity
BF1: bite force in the field
BF2: bite force in captivity
HL: head length
HH: head height

#### Work

Residual bite forces (from regression of log bite force on log mass) were plotted against duration of digging. No relationship was found between total work and mass-corrected bite force (Fig. 10.6, r= −0.3219;p= 0.19). Work is thus a poor predictor of biting performance.

**Fig. 10.6.**
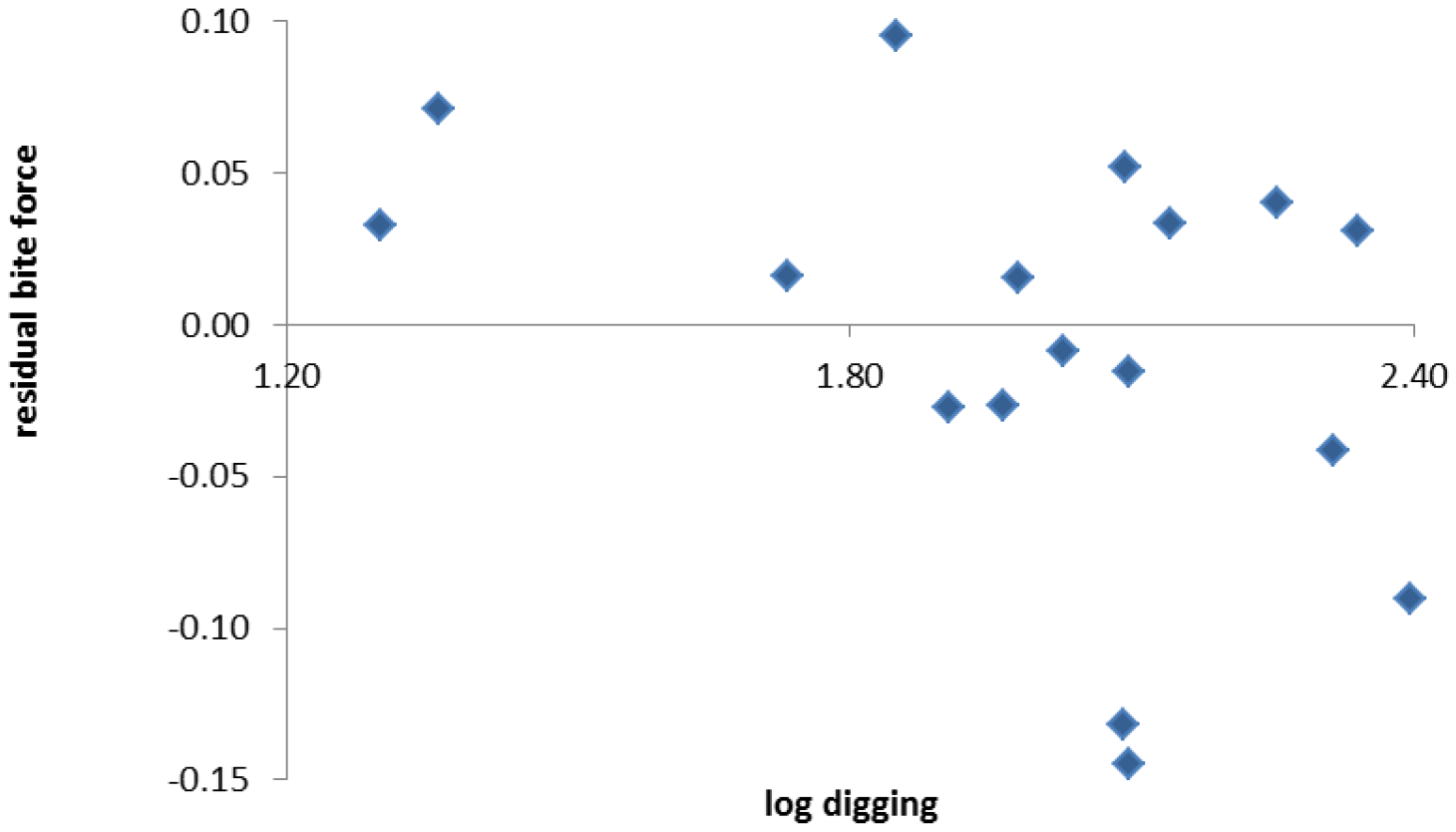
Residual from regression analysis of log10 bite force on log10 mass plotted against log10 of durations of digging behaviour.

#### Morphology

In the study colony, bite force was highly correlated with body mass (r= 0.97; p < 0.001) for measurements in the field and for measurements in captivity (r= 0.96; p = 0.001); Fig.10.7). Body mass is thus a good predictor of bite force. The correlation between log head height and log bite force for measurements in the field (r= 0.74; p = 0.0003) and for measurements in captivity (r= 0.77; P = 0.0002) was stronger than the correlation between log head length and log bite force for measurements in the field (r= 0.63; p = 0.004) and for measurements in captivity (r= 0.61; p = 0.007) (Fig. 10.8).

**Fig. 10.7.**
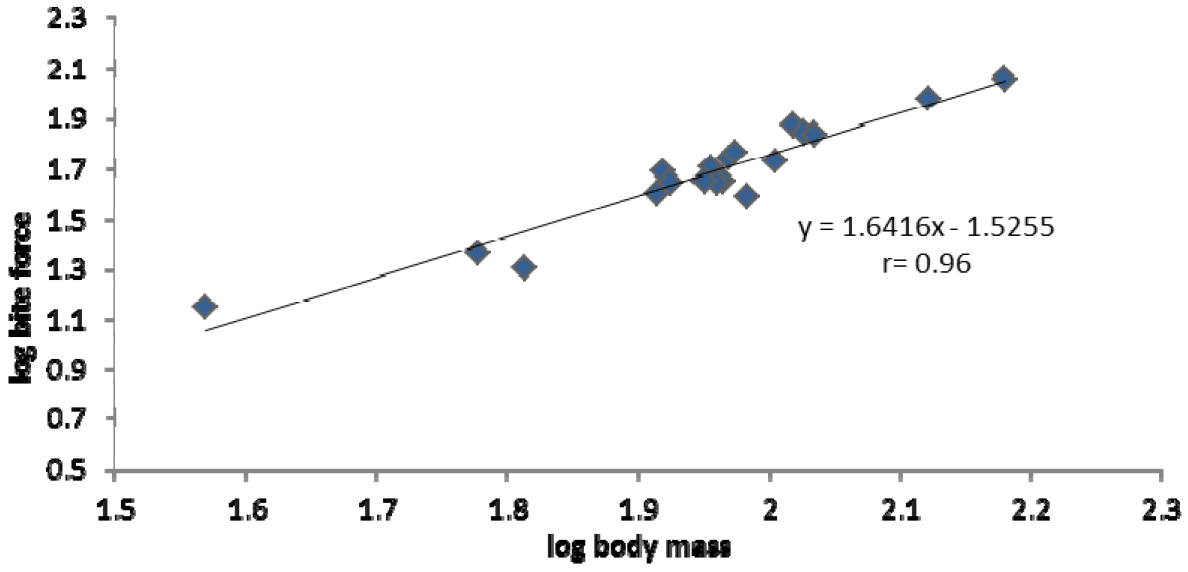
Relationship between bite force and body mass for colony of *Fukomys micklemi*.

**Fig. 10.8.**
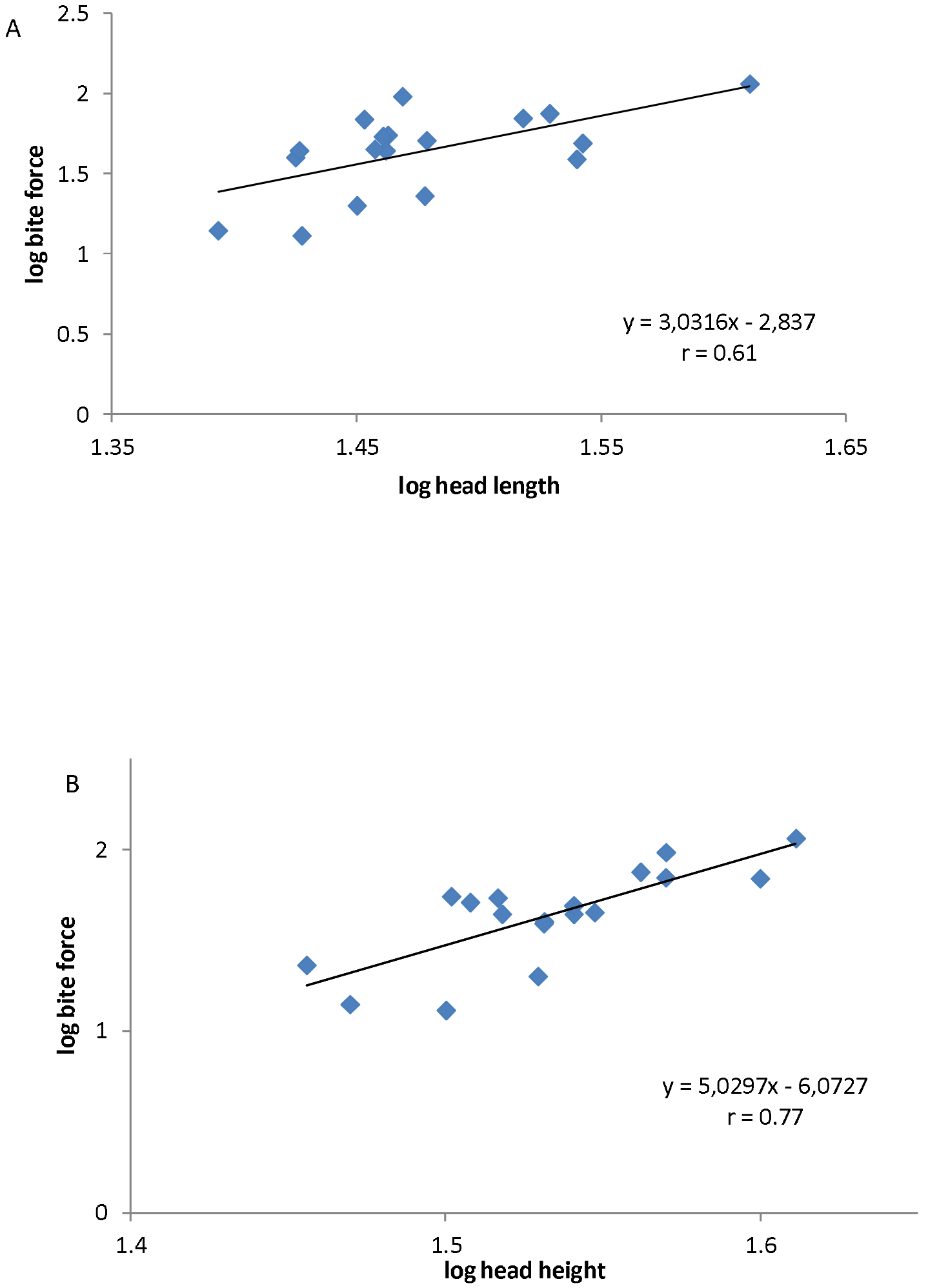
Relationship between bite force and A. head length B. head height for measurements of bite force in captivity.

## Discussion

### Castes

Based on the amount of work, no clear distinction could be made between a frequent and an infrequent worker caste. The existence of worker castes was first shown for naked mole-rats by Jarvis (1981). She made a distinction between frequent workers, infrequent workers and non-workers. Further research on naked mole-rats, however, has shown variation in the amount of work, but no distinct castes could be identified either (Lacey and Sherman, 1991; Bennett and Faulkes, 2000). The existence of different worker castes has also been described for other bathyergids such as *Cryptomys hottentotus* (Bennett, 1989; Moolman et al., 1998), *Fukomys* mechowii (Wallace and Bennett, 1998) and *Fukomys damarensis* (Bennett and Jarvis, 1988; Bennett, 1990; Jacobs et al., 1991). However, these studies don’t always show a clear gap in the amount of work. Wallace and Bennett (1998) define frequent workers as those performing 15-22% of the work and infrequent workers as individuals performing 7-11% of the work. Moolman et al. (1998) even describe three worker castes – infrequent workers, intermediate workers and frequent workers – for *Cryptomys hottentotus*. These studies concluded that *Fukomys* mechowii displays an intermediate degree of sociality between that of *Cryptomys hottentotus* and the two eusocial species (the naked and Damaraland mole-rat). Finally, although worker castes seem to be more distinct in *Fukomys damarensis*, for one colony Jacobs et al. (1991) use a difference of only 1% as a cut-off value to distinguish between frequent and infrequent workers. The amount of work performed by individuals within that colony thus rather seems to show a continuous distribution. Jarvis and Bennett (1993) used the term ‘loosely defined working groups’. We argue that the distinction between frequent and infrequent worker castes is a subjective one and does not reflect the dynamic pattern of development and behaviour in mole-rats (see below). There is considerable variation in work and different individuals might perform slightly different roles in the colony (for example ID 33 which could be called an allogroomer). However, we are far from a clear distinction between irreversible, permanent behavioural castes as can be found in eusocial insects.

It could be argued that focusing on work behaviour ignores other important behaviours, e.g. colony defence. Such behaviour is likely to influence the designation of individuals to a worker caste. Although clear patroller castes, soldier castes and worker castes will not likely be clearly distinguishable in *Fukomys*, defence – type behaviours for example may need further attention pending a final evaluation. Similarally required are studies looking caste-ralted physiological differences. O’Riain et al (1996) present data, which indicate a physiologically distinct dispersing caste in *Heterocephalus*. These individuals have more fat reserves and increase their energy expenditure after rainfall, whereas frequent workers do not. The frequent and infrequent workers were distinguished using morphological differences, based on the finding that infrequent workers have larger heads and longer bodies than frequent workers (Bennett and Jarvis, 1988; Bennett and Faulkes, 2000). Combining information from morphology, physiology and behaviour may lead to the identification of individuals with more or less specific roles within the colony.

The results confirm some of the findings from previous studies on the social structure of mole-rat colonies. Males and females did not differ in the amount of work. Bennett (1990) found that both sexes were represented in both the frequent and infrequent worker caste. Similarly, Wallace and Bennett (1998) found that there were no significant differences in the amount of work performed by each sex. The fact that breeding animals perform little work, confirms another common pattern among mole-rat colonies (Bennett 1989; 1990; Wallace and Bennett 1998).

Studies on naked mole-rats initially stated that the frequent worker caste consisted of the smaller and lighter individuals of the colony (Jarvis, 1981). The infrequent workers were larger and heavier. Later studies on naked-mole rats, however, did not always support these findings (Lacey and Sherman, 1991). Studies on *Fukomys* mechowii (Wallace and Bennett, 1998) and *Fukomys damarensis* (Jacobs et al., 1991) also did not support these findings. Bennett (1990), however, concludes that the castes differed significantly in body mass. Our results provide only mild support for the findings of Jarvis (1981). While there is a slight tendency for smaller individuals to perform more work, this is not a consistent pattern.

Bennett (1990) and Burda (1990) found that age polyethism is apparent in respectively *Fukomys damarensis* and *F. anselli*. Polyethism is defined as a functional specialization within a colony of social animals, leading to a division of labour. Age polyethism hereby refers to individuals changing castes as they grow older. This is in agreement with the situation for naked mole-rats (Jarvis, 1981). The young recruited to the colony join the worker caste. While the faster growing individuals quickly become infrequent workers, the slower growing animals remain in the worker caste. Lacey and Sherman (1991) argue that there is no real age or size polyethism, which is limited to a certain caste. In their view, the polyethism represents a flexible (albeit predictable) series of behavioural changes, linked to ageing and/or growth of an individual. The pattern of behaviourally development of a certain individual will be influenced by ecological and demographic factors (for example dominance). Jarvis et al. (1991) state that applying the term ‘age polyethism’ is an oversimplification of the ontogeny of a mole-rats behaviour. Consequently, differences in work behaviour between large and small or old and young individuals will be gradual and continuous. Although the youngest individual in the study colony (ID 63) performed the second largest amount of the work, other individuals, presumably young individuals (based on body mass), performed much less work. Thus, our results do not provide clear support for age polyethism in *F. micklemi*.

Accurate description of age and size polyethism, require monitoring of growth and development in captive born individuals. Furthermore, factors such as dominance structure should be taken into account. Jarvis et al. (1991) conducted an extensive study on naked mole-rats focusing on developmental patterns. A similar study has been done by Bennett for *Cryptomys hottentotus* (1989) and *Fukomys damarensis* (1988). However, further work is needed to assess possible differences in colony development between species taking different positions in the sociality continuum (Wallace and Bennett, 1998). It is possible that in eusocial species growth rates will show more variation between subsequent litters. Jarvis et al. (1991) hypothesized that the first litters will grow fast in order to quickly create a workforce. Subsequent litters may show slower growth rates to reduce energy costs of the colony. In colonies with less tight social structures these differences may be largely absent, since there may be an emphasis on fast growth to quickly gain a large size, which aids dispersal (Hazell et al., 2000; Spinks et al., 2000; Molteno and Bennett, 2002).

If differences in the pattern of development along the sociality continuum are apparent, this would allow testing predictions made by the AFDH. According to the AFDH, colonies inhabiting arid regions will have a better defined social structure than colonies inhabiting mesic regions (Faulkes and Bennett, 2007). Ideally, different species inhabiting the same ecotype could be used for defining developmental differences along the sociality gradient. Subsequently, colonies of a single species (or chromosome race) inhabiting different ecotypes could be used to test whether colonies from mesic habitats show similarities in development with species that display a lower degree of sociality and colonies from arid habitats show similarities with eusocial species.

Juveniles may be recruited to perform a lot of the work (c. q. into the frequent worker caste). As an individual grows, the animal would then gradually reduce its work input. This could be dependent on colony size, dominance of the individual and/or the presence of new recruits. The age or body mass at which an individual becomes an infrequent worker may thus show considerable variation. Failure to find distinct castes could be the result of a continuous process in which individuals are gradually shifting from frequent to infrequent worker. When observations are being carried out in stable periods when no caste-shifts are being made, clear-cut worker groups may appear. These groups might then disappear when some individuals are making the transition from frequent to infrequent worker. Furthermore, there might be differences in the general work package of different individuals (i.e. type and amount of work, see above). Long-term studies on a individual colonies could clarify whether such a dynamic model exists.

Results from this study show that, although there is considerable variation in work, *Fukomys micklemi* does not appear to show a well-defined subdivision of the non-reproductive caste. Morphological variation could not be incontestably linked to the amount of work. Thus, no evidence for the existence of clearly distinct morphological and/or behavioural castes was found.

### Biting performance

#### Field versus captivity

The data indicate that mole-rats in captivity will show an increase in mass, which is independent of growth. If the increase in body mass would be the result of growth, then no decrease in bite force should be expected, since bite force changes linearly with body mass and head measurements. The increase in body mass is most likely caused by an increase in fat tissue. As a result of the increase in fat reserves, the body mass increases relatively faster than the muscle mass, resulting in a decrease of bite force per unit of mass.

#### Work

No relation was found neither between bite force and work, nor between mass-corrected bite force and work. It could be hypothesized that individuals that bite harder do so because they have larger jaw muscles and a coinciding higher body mass. Similar results were found when correcting for head length or head height. The relation between bite force and work, therefore remains unproven.

#### Morphology

Bite force was highly positively correlated with body mass, indicating that body mass is a very good predictor of bite force. Confirming results from Van Daele et al (2009), bite force was more strongly correlated with head height than with head length. Therefore head height is a better predictor for bite force than head length. The strongest correlation was found between bite force and body mass.

The results from this study show that there is a large amount of variation in morphological parameters such as body mass, head length and head height within a single colony of *Fukomys micklemi*, as well as a substantial amount of variation in biting performance. It is clear that the morphological parameters are intricately linked to biting performance and will generally be good predictors of bite force. There is also marked variation in the amount of work a certain individual will perform. This behavioural variation, however, is correlated with neither the morphological variables nor biting performance. Thus, it seems that biting performance is not the result of differential selection pressure on animals that differ in the amount of work.

Preliminary results from studies on other *F. micklemi* colonies indicate that sexual differences in biting performance may occur and interact with the influence of captivity (own data; not shown). Sexual difference may be linked to dominance hierarchy, a factor which has not been studied in *F. micklemi* to date. It has indeed been shown for lizards that sexual differences in biting performance are related to dominance (Huyghe et al., 2005). Jacobs et al. (1991) found that males of *Fukomys damarensis* were on average more dominant than females. If this would be consistent across colonies of *Fukomys micklemi*, this would allow testing whether these sexual differences are a result of the dominance structure.

It can be expected that ecological factors will play an important role in the fitness effect of biting performance. The studied specimens came from the more arid part in the species’ distribution, characterized by hard soils during the dry season. This could have implications for biting performance. It could thus be hypothesized that colonies in arid habitats achieve high bite forces, which would be adaptive for digging in hard soil. However most of the digging is done during the rainy season, when the soil is softer (Jarvis et al., 1998). On the other hand, arid habitats are expected to put energetic constraints on colonies (as predicted by the AFDH). Consequently, bite forces may not be primarily determined by work. Variation in biting performance may be the result of variation in developmental patterns (see above).

This was a pioneering study concerning the link between bite force and (eu)sociality in mole-rats and the first study to assess patterns of work for *Fukomys micklemi*. Further research is needed, whereby more colonies of *Fukomys micklemi* from arid as well as mesic habitats are studied. Morphological, physiological and behavioural data should be combined and linked to developmental patterns. Furthermore, differences between dry season en rainy season should be assessed, as this may be another factor influencing social structure.

